# ProDCoNN-server: a web server for protein sequence prediction and design from a three-dimensional structure

**DOI:** 10.1101/2021.11.04.467289

**Authors:** Yuan Zhang, Arunima Mandal, Kevin Cui, Xiuwen Liu, Jinfeng Zhang

**Affiliations:** Department of Statistics, Florida State University, Tallahassee, FL 32306; Department of Computer Science, Florida State University, Tallahassee, FL 32306; Department of Computer and Information Science and Engineering, University of Florida, Gainesville, FL 32611

## Abstract

We present ProDCoNN-server, a web server for protein sequence design and prediction from a given protein structure. The server is based on a previously developed deep learning model for protein design, ProDCoNN, which achieved state-of-the-art performance when tested on large numbers of test proteins and benchmark datasets. The prediction is very fast compared with other protein sequence prediction servers - it takes only a few minutes for a query protein on average. Two models could be selected for different purposes: BBO for full sequence prediction, extendable for multiple sequence generation, and BBS for single position prediction with the type of other residues known. ProDCoNN-server outputs the predicted sequence and the probability matrix for each amino acid at each predicted residue. The probability matrix can also be visualized as a sequence logos figure (BBO) or probability distribution plot (BBS). The server is available at: https://prodconn.stat.fsu.edu/.

## Introduction

Designing protein sequences that fold to a given three-dimensional structure, known as inverse protein folding (IPF), has long been challenging in computational structural biology with significant theoretical and practical implications. Solving this problem will improve our fundamental understanding of the sequence-structure relationship of proteins. There have been some significant successes in IPF in the past. The traditional methods are usually categorized into two approaches. One is an energy-based method, which starts with random protein sequences and iteratively optimizes an energy function via mutations until the energy score reaches a minimum ^1–6^. The other one is using local fragment structures from a target structure, which is compared to the fragment library of known protein structures ^7–10^. The traditional methods are either time-consuming or rely on the availability of structures in the fragment library.

In recent years, deep learning methods based on neural networks have dramatically impacted the computational biophysics field, which helps to solve IPF problems. The SPIN^11^ (Sequence Profiles by Integrated Neural network) based on fragment-derived sequence profiles and structure-derived energy profiles yielded an average sequence recovery of 30.7% for a dataset with 1532 proteins. Later, SPIN was upgraded to SPIN2^12^ and achieved an average sequence recovery of 34.4%. Another study adopting a DNN for protein design, conducted by Wang et al.^13^ in 2018, used structure features as input and yielded the best recovery rate at 34% on a dataset with 10173 proteins (30% sequence identity). We developed ProDCoNN^14^ based on a convolutional neural network (CNN) to predict the residue type along the sequence. The model took a gridded box with the atomic coordinates and types around a residue as input. ProDCoNN achieved an accuracy of 42.2% for the test dataset (30% sequence similarity). Later, Qi et al. designed DenseCPD^15^ using a DenseNet architecture and improved accuracy to 50.96%. However, because the mapping from sequence to structure is not unique, it is not clear that higher sequence recovery rates would be meaningful.

Here, we present ProDCoNN_server, a web server for protein sequence prediction and design. The server takes a three-dimensional structure (pdb format) as input and outputs the predicted sequence and the predicted probabilities of 20 amino acids for each residue. Since it is well-known that proteins can tolerate mutations on most of their sequence positions, the probability matrix could give more information, which could be used for an in-deep analysis and sequence design. A logo figure^16^ (BBO model) based on the probabilities, or a probability distribution plot (BBS model) will be generated. The prediction on our server is in no time, and one job could be finished within a few minutes, which is much faster than other protein sequence prediction servers, such as RosettaDesign server^17^ and DenseCPD server.

## Method

### 1. ProDCoNN

ProDCoNN tackles the protein design problem by predicting one residue at a time, called target residue, using the local structural information surrounding the target residue. A gridded box centered on the Cα atom of the target residue is used to capture the local structural information. The edge of the gridded box is 18 Å, with each voxel being unit size (1Å × 1Å × 1Å). We use three-dimensional truncated Gaussian functions to smooth the input data to overcome the limitation caused by the discretization of the 3D space around the target residue. The information will be sent to a pre-trained model for prediction. We have two models for different applications: Backbone only model (BBO) takes protein backbone conformation information as input which is suitable for full sequence prediction beginning with backbone structures only. Backbone with sequence model (BBS) takes backbone information plus C_β_ atoms of non-target residues labeled as one of the 20 amino acid types based on the sequence information. This model requires sequence information except for target residue, which is suitable for predicting a single residue given the backbone structure and the amino acid types of the rest of the sequence. Our server only uses the BBO_ID90 and BBS_ID90 models of ProDCoNN, trained by using a dataset with 21,071 protein structures from PDB with sequence identity lower than 90%.

### 2. Input

ProDCoNN_server takes a PDB structure, in pdb format, as input for prediction, which should be uploaded at the “Upload File” section. The PDB file must follow the standard pdb file format. Columns 18-21 should not be empty, which could be ‘ALA’ if the residue type is unknown.

In the “Filter” section, the client must specify the chain name, as a single alphabet, for prediction. And the prediction range could be selected for partial sequence prediction. We use residue index (count from 1 from the first residue in each chain in the pdb file) instead of residue sequence number (column 23-26 in the pdb file) to define the prediction range. The default value is “1” for the beginning and “-1” for the end, which will predict every residue in the chain. If using the BBS model, both the beginning and end should be set as the index of the predicted residue.

In the “model settings” section, model type should be selected to make the full sequence prediction (BBO) or single residue prediction (BBS). The default value is “BBO”. In the “email results” section, an email is required to get the resultant text file (predicted sequence with the probability matrix) via email.

### 3. Output

After submitting the job, the client can either wait for the result on the server’s result page or close the page, and the results will be sent to the registered email. The output page, as shown in Figure 1, displays the input sequence (the sequence in the input pdb file) and the predicted sequence by our model. The predicted probability matrix in a text file, which contains the probabilities of 20 amino acids for each predicted residue, could be downloaded. In the probability matrix file, the first column is the input amino acid. The second column is the predicted amino acid, then follows the probabilities for amino acids A, C, D, E, F, G, H, I, K, L, M, N, P, Q, R, S, T, V, W, and Y.

**Figure 1:**
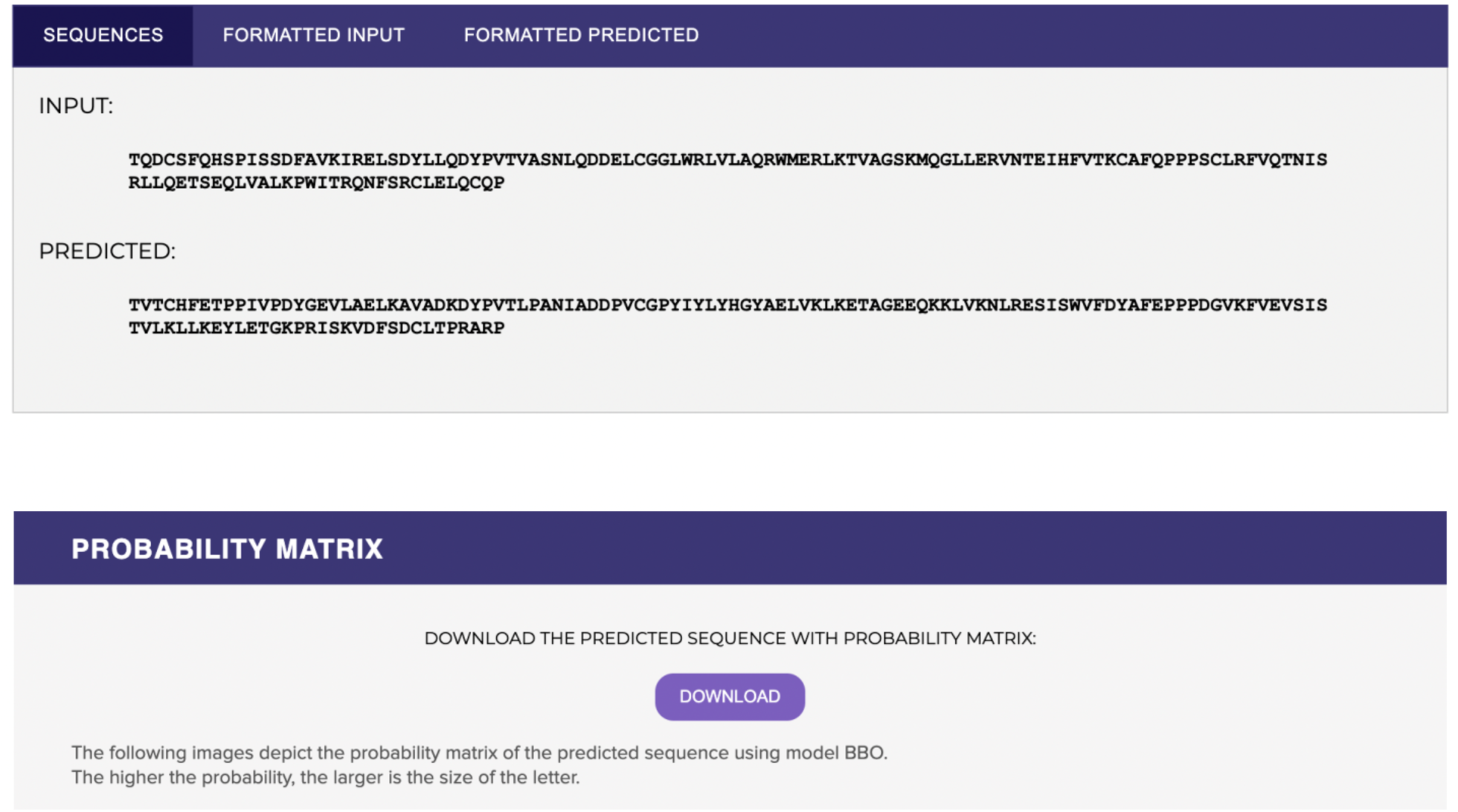
The ProDCoNN_server output interface.

A sequence logos figure (Figure 2 (a)) is generated if the BBO model is used, which shows the top 6 predictions of each residue along the sequence. A figure (Figure 2 (b)) shows the probabilities of 20 amino acids for the target residue if the BBS model is used.

**Figure 2:**
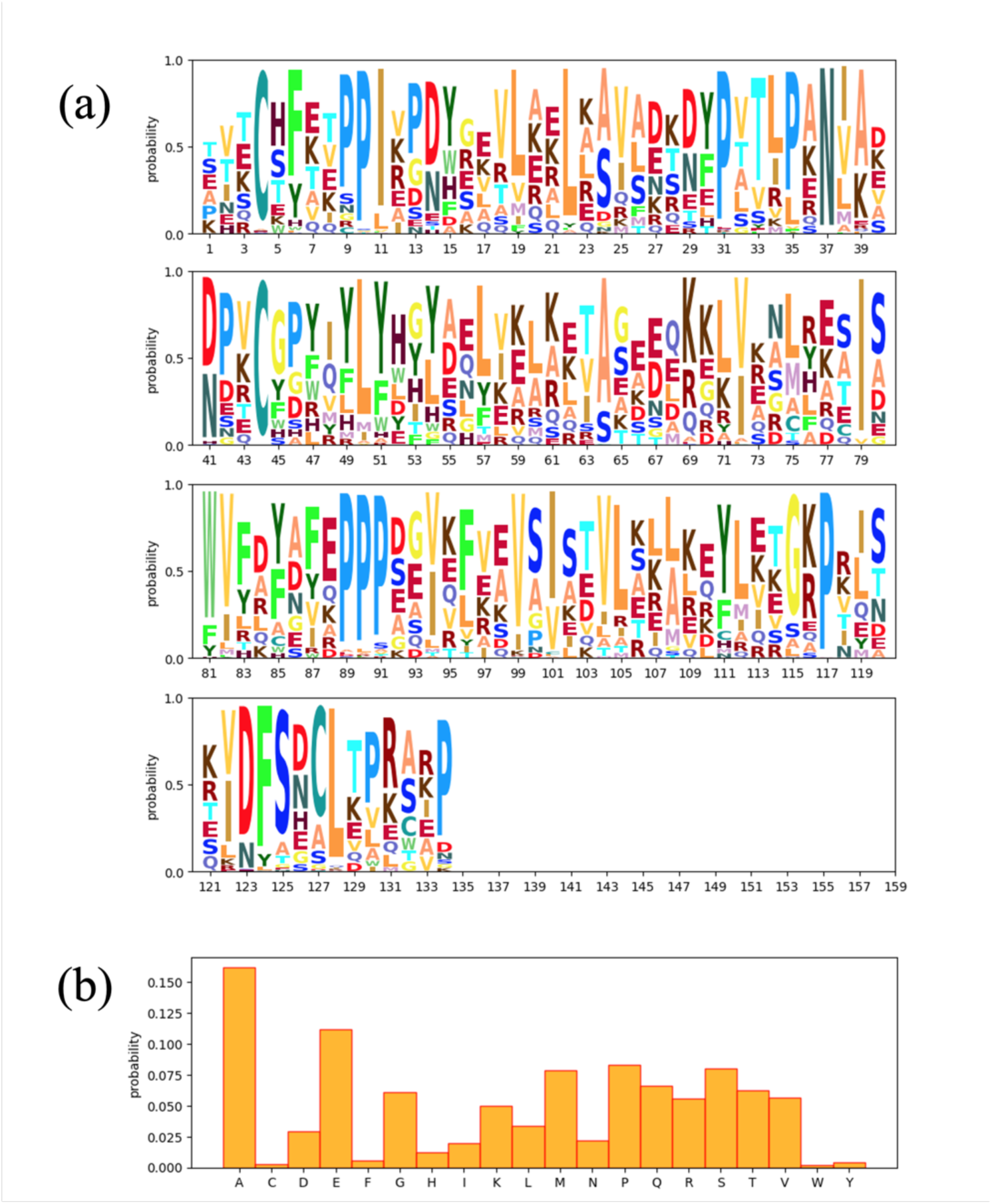
Visualization of the result for BBO (a) or BBS (b) model, which is shown at the output interface.

## Summary

ProDCoNN_server provides a platform to perform protein sequence prediction from an uploaded pdb structure. Two models could be selected for different purposes: BBO for full sequence prediction, which could be extended for multiple sequence generation, and BBS for single position prediction with other residue types known, which is helpful for some applications like single mutation experiments. Both models will output the predicted sequence and the probability matrix for each amino acid at each predicted residue. And the probability matrix is also visualized as a sequence logos figure (BBO) or probability distribution plot (BBS). The prediction on our server is very fast, and one job could be finished within a few minutes, which is much faster than other protein sequence prediction servers, such as the RosettaDesign server and DenseCPD.

We plan to add a new partial-labeled model (PBS) and implement a sequential Monte Carlo method to do protein sequence sampling from a given pdb structure and estimate the designability of the structure {cite}.

## Supported platforms

All latest web browsers are supported.

## ACKNOWLEDGMENTS

Research reported in this publication was supported by the National Institute of General Medical Sciences of the National Institute of Health under award number R01GM126558. The content is solely the responsibility of the authors and does not necessarily represent the official views of the National Institutes of Health.

## Notes

https://prodconn.stat.fsu.edu/

### Competing Interest Statement

The authors have declared no competing interest.

